# RepeatProfiler: a pipeline for visualization and comparative analysis of repetitive DNA profiles

**DOI:** 10.1101/2020.05.22.111252

**Authors:** S. Negm, A. Greenberg, A.M. Larracuente, J.S. Sproul

## Abstract

Study of DNA repeats in model organisms highlights the role of repetitive DNA in many processes that drive genome evolution and phenotypic change. Because repetitive DNA is much more dynamic than single-copy DNA, repetitive sequences can reveal signals of evolutionary history over short time scales that may not be evident in sequences from slower-evolving genomic regions. Many tools for studying repeats are directed toward organisms with existing genomic resources, including genome assemblies and repeat libraries. However, signals in repeat variation may prove especially valuable in disentangling evolutionary histories in diverse non-model groups, for which genomic resources are limited. Here we introduce *RepeatProfiler*, a tool for generating, visualizing, and comparing repetitive DNA profiles from low-coverage, short-read sequence data. *RepeatProfiler* automates the generation and visualization of repetitive DNA coverage depth profiles and allows for statistical comparison of profile shape across samples. In addition, *RepeatProfiler* facilitates comparison of profiles by extracting signal from sequence variants across profiles which can then be analyzed as molecular morphological characters using phylogenetic analysis. We validate *RepeatProfiler* with data sets from ground beetles (*Bembidion*), flies (*Drosophila*), and tomatoes (*Solanum*). We highlight the potential of repetitive DNA profiles as a high-resolution data source for studies in species delimitation, comparative genomics, and repeat biology.

## Introduction

Repeats have been understudied for decades due to technical and computational challenges associated with their sequencing and assembly. As recent advances in sequencing technology overcome those obstacles, work in model organisms is shedding new light on the critical roles repeats play in many processes including shifts in gene regulation that drive phenotypic change, rapid genome evolution, and mechanisms underlying reproductive isolation and speciation (Chuong, Elde, & Feschotte, 2017; Ferree & Barbash, 2009; Niu et al., 2019; Schrader & Schmitz, 2019; Stuart et al., 2016). Because repetitive regions can evolve much more rapidly than unique DNA sequences (*e.g.*, protein-coding genes), repetitive sequences can reveal signals of evolutionary history over short time scales that may not be evident in slower-evolving genomic regions.

Despite their critical roles in genome and phenotype evolution and their known rapid turnover between closely related species (*e.g*. (Lower et al., 2017; Sproul, Khost, et al., 2020; Strachan, Coen, Webb, & Dover, 1982; Tetreault & Ungerer, 2016; Ugarković & Plohl, 2002)), repeat dynamics are seldom considered in evolutionary studies aiming to understand species boundaries and recent genome evolution. One caveat to using repeats in evolutionary studies is that their abundance may fluctuate widely across samples, even below the species level (Bosco, Campbell, Leiva-Neto, & Markow, 2007; McLain, Rai, & Fraser, 1987; Mestrović, Plohl, Mravinac, & Ugarković, 1998; Raskina, Barber, Nevo, & Belyayev, 2008). Within species, this variation may be largely caused by expansion and contraction of repeats due to unequal exchange of recombining repeat clusters (Smith, 1976). Dynamic variation in repeat abundance may add noise when comparing patterns across samples, which can confound signals that may be present (Martín-Peciña, Ruiz-Ruano, Camacho, & Dodsworth, 2019). Development of approaches that can extract evolutionary signal despite repeat abundance variation can potentially increase the resolution of evolutionary studies among populations and species (Sproul, Barton, & Maddison, 2020).

Many software tools are available for studying repetitive DNAs. Most tools rely on fully assembled genomes and repeat libraries (Goerner-Potvin & Bourque, 2018) and are therefore most useful in groups with extensive genomic resources. However, a number of “reference-free” tools enable discovery and annotation of repeats in any group using low-coverage shotgun sequence data (Ewing, 2015; Nelson, Linheiro, & Bergman, 2017). Despite these advances, a general shortcoming of repeat software is that few programs have features that allow for quantitative comparison of repeat patterns across samples. Such comparative analysis will become increasingly important as the study of repeats extends from few representative individuals to more extensive sampling (*e.g.*, population-level sampling within species), which is important for understanding how repeats shape genome evolution within and across species. In addition, repeats present a special set of challenges to standard comparative genomics approaches (Fernandes et al., 2020). Repeats account for large fractions of missing sequence in assemblies—this remains true even in premier model organisms with ‘gold standard’ genomic resources. It is difficult to conduct any comprehensive study of repeats on incomplete assemblies. Although long-read technology is closing many of these gaps, the megabase-long blocks of tandem repeats that are common in many genomes remain a challenge to assemble for the foreseeable future. Furthermore, reads arising from repetitive sequences present multiple mapping issues (*i*.*e.*, one read can map equally well to multiple places in the genome) adding uncertainty to analyses that rely on mapping reads to assembled repeats. Approaches that measure repeat dynamics directly from low-coverage sequence data have advantages for detecting signatures of repeat dynamics that may not be evident in incomplete genome assemblies and for studies in groups with limited genomic resources.

Here we present *RepeatProfiler* as a tool for both exploration and comparative analysis of repetitive DNA patterns with low-coverage, short-read data. The pipeline was motivated by a previous study by Sproul et al. (2020), which highlighted the potential of repetitive DNA profiles in species delimitation studies. *RepeatProfiler* maps reads to reference repeat sequences, generates enhanced read depth/copy number profiles for visualizing patterns, and facilitates statistical comparison of profiles within and among samples. The pipeline compares sources of profile variation (*e.g.*, profile shape, and relative abundance of variants within profiles) that can be stable within species despite variation in absolute repeat abundance. Comparing profiles for the same repeat reference between multiple samples can reveal species-specific differences in sequence composition (*e.g.*, substitutions, indels), relative abundance, truncations, and amplification of partial repeats throughout the genome. This approach to studying repeats bypasses issues related to multiple mapping and incomplete genome assemblies because all genomic reads are mapped to reference sequences and the resulting profiles capture variation in the repeat sequence, regardless of their genomic distribution (Fernandes et al., 2020; Sproul, Barton, et al., 2020). Studying profiles of multiple repeats in a comparative framework can provide evidence of gene flow boundaries, highlight potential mechanisms underlying repeat evolution, and detect signatures of rapid genome evolution not evident in less dynamic components of the genome (Sproul, Barton, et al., 2020; Sproul & Maddison, 2018).

*RepeatProfiler* uses Bowtie2 (Langmead & Salzberg, 2012), SAMtools (Li et al., 2009), and python to map reads and process mapping output. It uses R (Team, 2013) including the packages ggplot2 (Wickham, 2016a), ggpubr (Kassambara, 2017), reshape2 (Wickham, 2007), and scales (Wickham, 2016b) for data visualization and comparative analysis of profile features. The pipeline itself is written in bash and published under GPL 3.0 license. *RepeatProfiler* is available for download from https://github.com/johnssproul/RepeatProfiler and can be installed on Unix, Linux, and Windows platforms with installation options through Homebrew and Docker to automate installation of dependencies.

## Materials and Methods

### Overview

An overview of the pipeline workflow is provided in Figure S1. *RepeatProfiler* generates repetitive DNA profiles by mapping input reads to reference sequences of repeats of interest. Output includes two styles of enhanced read depth profiles (color-enhanced and variant-enhanced) to facilitate comparison of patterns across samples and repeats. Output also includes summary tables and results from comparative analysis.

### Input

*RepeatProfiler* has two input requirements: (1) short-read sequence data from one or more samples in FASTQ format; (2) a FASTA file containing reference sequences of repeats to be analyzed. Short-read sequence data may include low-coverage (*e.g*., 0.1–10X) whole-genome shotgun data or reads produced from a target-capture sequencing approach (*e.g*., anchored hybrid enrichment or ultra-conserved element approach). The latter assumes input reads contain off-target ‘background’ reads such that profiles can be generated from off-target reads while ignoring reads from enriched targets. Repeat reference sequences may be obtained from existing repeat libraries or online databases (*e.g*., NCBI, Dfam). For organisms lacking reference libraries and/or representation in databases, reference sequences may be obtained using repeat assembly software such as RepeatExplorer (Novák, Neumann, Pech, Steinhaisl, & Macas, 2013) with possible workflows outlined in Supplemental Material. The pipeline assumes input reads have undergone quality control (*e.g*., trimming and adapter removal) and downsampling (if desired) prior to analysis.

### Generating profiles

The pipeline generates read depth profiles by mapping input reads to reference sequences using Bowtie2. We use Bowtie2 as it has been shown using real data to have higher rates of read mapping while still being faster than comparable programs (Langmead & Salzberg, 2012). *RepeatProfiler’*s default settings use Bowtie2 parameters that tolerate mismatches, given that users may be mapping reads from divergent species to a common reference; however, users can alter read mapping parameters for other uses of the tool. Following read mapping, bam files are processed using SAMtools, including the mpileup function which generates variant information at each site relative to the reference sequence. SAMtools output is then parsed to retain read depth information and simplify variant output prior to plotting for visualization.

### Visualizing profiles

*RepeatProfiler* produces two types of enhanced read-depth profiles in R using the ‘ggplot2’, ‘ggpubr’, ‘scales’, and ‘reshape2’ packages. Color-enhanced profiles use a color gradient to provide a visual indication of read depth at each site using a scale that is standardized across samples and reference sequences. Variant-enhanced profiles provide a visual summary of sites that show sequence variants relative to the reference sequence. For both color and variant-enhanced profiles we combine plots into a single, multi-plot PDF to simplify visual comparison of patterns across samples and repeats.

### Comparative analysis

#### Correlation analysis

Using an optional flag (-corr), the user can test the degree of correlation in profile shape (*i.e.*, the pattern of coverage depth across a given reference sequence) within and among user-defined groups. Sample groups may represent different species, populations, sexes, tissues, or other groupings. The correlation analysis measures the degree of similarity in coverage depth for each position across the reference sequence between two samples. We use Spearman’s rank correlation coefficient (or Spearman’s rho, denoted ‘*ρ*’) to calculate correlation values for pairwise comparisons of all samples for a given reference sequence. Because repeat abundance is expected to vary even within populations, we use ranks instead of absolute coverage values for correlation analysis. Beyond population-level variation, variation in genome size, the number of input reads, and bias in library preparation/sequencing among samples can all lead to further variation in absolute repeat abundance. Thus, Spearman’s rho allows for comparison of profile shape while minimizing the effect of noise caused by actual, and introduced, repeat abundance differences across samples.

#### Phylogenetic analysis of variant signatures

*RepeatProfiler* also facilitates profile comparison in a phylogenetic framework. The pipeline summarizes information contained in variant-enhanced profiles by identifying abundant variants and encoding those variants as molecular-morphological characters. The pipeline outputs PHYLIP files with the encoded pattern of variants (relative to the reference sequence) which the user can then analyze as morphological data using phylogenetic software. If the user analyzes multiple repeats in the run, trees can be produced from the output of each repeat and summarized as a consensus tree to combine signal from all repeats into one tree. The intent behind this feature is not to infer phylogenetic relationships among samples *per se* (though our validation below suggests the pattern of variants can accurately resolve phylogenetic relationships over short times scales), but rather to take advantage of the statistical framework of phylogenetic analysis to group profiles that have similar variant signatures, and to indicate the strength of that grouping (as indicated by nodal support). Similar to the correlation analysis, this approach is robust to absolute differences in read depth as it relies on the relative abundance of variants within a sample, rather than absolute values which vary due to many factors described above. This feature may be used to test whether signal from profiles supports *a priori* grouping of samples or to generate grouping hypotheses in the absence of existing data. Additional details regarding the implementation of this feature are provided in Supplemental Material.

#### Single-copy normalization

As an optional feature, the pipeline conducts normalization of read depth across samples relative to single-copy genes. This feature is useful if the user desires to make inferences about relative repeat copy number differences within or across samples. By choosing the ‘-singlecopy’ flag, the pipeline maps reads to user provided single-copy genes, estimates their average coverage, and calculates a normalized value of repeat coverage across samples based on single-copy estimates. Bases near the edges of a reference sequence are expected to show reduced coverage as an artefact of the read mapping algorithm, thereby underestimating coverage of single-copy genes (Pflug, Holmes, Burrus, Johnston, & Maddison, 2019) which would lead to overestimation of repeat copy number. To address this problem, the pipeline calculates a corrected value of single-copy coverage by excluding data near reference sequence edges (length of excluded sequence=1/2 of read length at each end of the reference) prior to calculating average coverage. The values from the correlation analysis are unaffected by this normalization since that analysis only considers ranks of coverage at each site.

### Validation methods

We validated the pipeline using previously published short-read data from ground beetles, tomatoes, and *Drosophila*, and simulated data from *Drosophila*. *RepeatProfiler* automates a general workflow of using repeat profiles in evolutionary studies presented in Sproul & Maddison (2018) and Sproul et al. (2020). Those studies explored reference bias, profile stability across varying read depth, tested whether profiles could be obtained from targeted sequencing workflows (*i.e.*, hybrid capture) and museum specimens, and used fluorescence *in situ* hybridization to relate information in profiles back to repeat architecture on chromosomes. Our validation here complements and extends those findings. A list of data sets used for the additional validation is provided in Supplementary Table S1.

### Read mapping parameter sensitivity analysis and normalization estimates

#### Parameter sensitivity analysis

We analyzed the effects of changing Bowtie2 parameters to determine ideal default settings for *RepeatProfiler* and to orient users as to the impact of parameter settings on resulting profiles. We generated profiles using Bowtie2’s presets: ‘very-fast’, ‘fast’, ‘sensitive’, and ‘very-sensitive’ with ‘end-to-end’ (Bowtie 2’s default) and ‘local’ mapping strategies. We conducted this analysis in a ground beetle species, *Bembidion breve*, using the 28S ribosomal RNA gene as a reference sequence. This species has a recent history of ribosomal DNA (rDNA) mobilization in which a fragment of 28S rDNA escaped functional rDNA units, proliferated, and spread throughout the genome where it now evolves separately (*i.e*., not in concert with functional rDNA) (Sproul, Barton, et al., 2020). Thus, this species and reference sequence pair is ideal for understanding the effect of changing mapping parameters on both highly conserved (*i.e.*, functional 28S rDNA) and more divergent (abundant rDNA-like fragments) reads relative to the same reference sequence.

#### Normalization estimates

An advantage of *RepeatProfiler’*s approach to studying repeat dynamics is that informative profiles can be obtained with very low-coverage sequence data (*e.g.*, much less than 1X coverage for abundant repeats). Because normalization relies on mapping reads to single-copy genes, we tested whether accurate single-copy estimates could be obtained with low-coverage data by running the pipeline across a series of downsampled data sets ranging from 25 million to 0.5 million reads (or 4X to ~0.1X coverage). We generated downsampled data sets using seqtk (https://github.com/lh3/seqtk) and mapped reads to ten, single-copy genes in two size classes (*i.e.*, 450-bp and 900-bp references) using the pipeline. We plotted the coverage estimates produced by each set of input reads to evaluate the stability of the estimate trend line at low coverage using R.

### Read mapping and reference sequence divergence

We investigated the effect of sequence divergence on read mapping rates by simulating divergent reference sequences in four closely-related *Drosophila* species (*D. melanogaster*, *D. imulans*, *D. sechellia*, and *D. mauritiana*) from species-specific reference sequences. We used SLiM 2 v.3.4 (Haller & Messer, 2017) to simulate sequence divergence over 2000 generations. We generated a sequence every 200 generations resulting in a total of ten new sequences with increasing sequence divergence such that the tenth showed ~25% pairwise divergence relative to the original sequence. We repeated simulations for two reference sequences (portions of the ribosomal DNA external transcribed spacer and the 28S ribosomal RNA gene) in each of the four species, ran the divergent reference sequences through the pipeline, and plotted the percentage of reads that mapped relative to the unmutated, conspecific reference sequence for each species.

### Phylogenetic validation

We validated our approach to condensing patterns in variant-profiles into molecular-morphological characters for phylogenetic analysis using empirical data from *Drosophila*, ground beetles, and tomatoes. We present a detailed validation using *Drosophila* because *D. melanogaster* and its close relatives are a group with: a very recent history of divergence (with known dates) (Garrigan et al., 2012), well-established phylogenetic relationships, abundant genomic resources (*i.e.*, sequence data and repeat libraries), and several repeats with a recent history of dynamic activity (Jagannathan, Warsinger-Pepe, Watase, & Yamashita, 2017; Larracuente, 2014; Lohe & Roberts, 1988; Sproul, Khost, et al., 2020). We obtained consensus sequences of 59 abundant transposable elements (TEs) including LTR and non-LTR retrotransposons from a custom *Drosophila* repeat library (Chakraborty et al., 2020; Sproul, Khost, et al., 2020) and generated profiles from these TEs in a run that included four closely related *Drosophila* species (*D. melanogaster*, *D. simulans, D. sechellia*, and outgroup *D. erecta*), with 2–4 replicate individuals from each species. The ingroup species diversified in the last 2.5 million years and two species (*D. simulans* and *D. sechellia*) are only separated by an estimated ~240k years (Garrigan et al., 2012). To reduce missing data in downstream analysis, we filtered run output to exclude any TEs with less than 70% average coverage of bases in the reference sequence for all samples. For the 37 remaining high-coverage TEs, we analyzed the PHYLIP file produced by the pipeline in IQ-TREE (Nguyen, Schmidt, von Haeseler, & Minh, 2015) as morphological data using an MK model. We processed resulting trees to test: (1) how many of the five expected clades are present with greater than 50% bootstrap support; (2) whether the branching pattern among those clades matches the species phylogeny; (3) whether a lack of expected clades is due to unresolved groupings of the closest sister pairs (*i.e.*, *D. simulans* + *D. sechellia*) or due to clades that group non-sister taxa; (4) whether any clades show >50% bootstrap support for relationships discordant with the species tree. In addition to analyzing individual gene trees we generated a consensus tree in IQ-TREE to evaluate the combined signal from all trees.

## Results

### Standard output

Enhanced repeat profiles produced by the pipeline (Figs. 1–2) can provide a high-resolution data source for evolutionary studies over short time scales. The characteristics of individual profiles can reveal species-specific signatures in both the pattern of coverage depth across the reference sequence (*i.e.*, profile shape) (Figs. 1A, S2) and the signature of sequence variants relative to the reference (Figs. 1B, S3–4).

**Figure 1.**
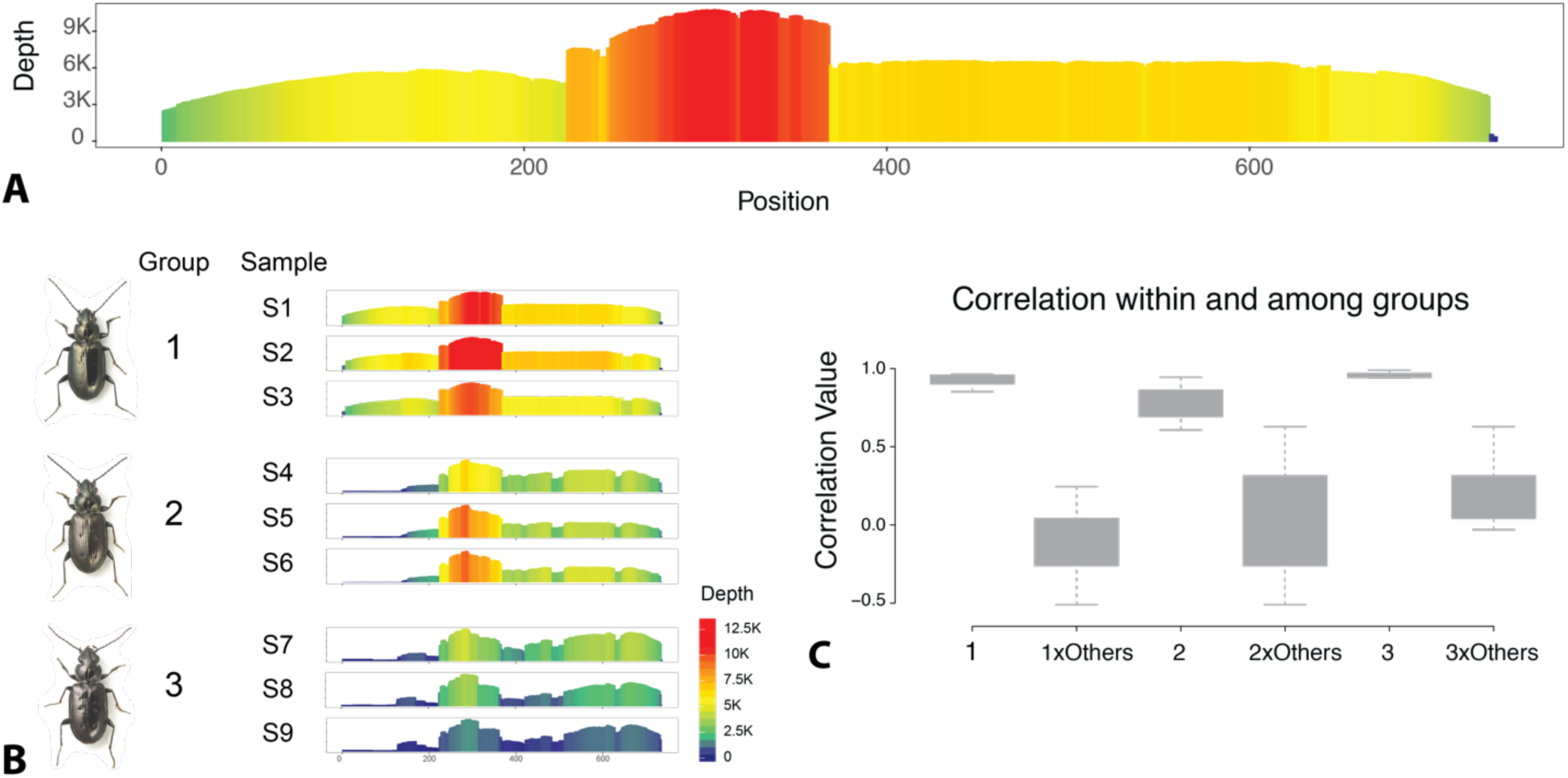
Examples of *RepeatProfiler* color-enhanced profiles and correlation analysis output. (A) A color-enhanced repeat profile from a putative satellite DNA in *Bembidion ampliatum* (Casey); color legend shown in B. (B) Profiles for the same satDNA in nine individual samples belonging to each of three closely related species: *B. ampliatum* (Group 1), *B. breve* (Motschulsky) (Group 2), and *B. lividulum* (Casey) (Group 3), with profiles showing species-specific signatures. (C) Output of correlation analysis with boxplots showing high correlation of profiles shape within groups, and low correlation between groups. Sample numbers, group numbers, and beetle images were added to the raw output of the pipeline for this figure.

In addition, profiles can show signatures that lend insight into repeat biology such as 5’ truncations in active non-LTR retroelements (Fig. S2) or male-female differences that can indicate differential distribution of repeats on sex chromosomes. The visual enhancements of color and variant-enhanced profiles simplify quick comparison across samples and repeats to understand general patterns while the comparative analysis features of the pipeline enable quantitative comparison for patterns of interest. Standard output also includes a table with summary statistics of coverage across samples and references.

### Comparative analysis of profiles

For multi-sample runs that include the correlation analysis flag (-corr), several additional output files are generated. For each reference sequence the program produces boxplots that summarize the distribution of pairwise correlation values (*i.e*., Spearman’s rho) for within and between-group comparison of profile shape (*e.g.*, Fig. 1C). An additional boxplot chart is produced that summarizes overall correlation patterns observed across references for each user group. Output also includes a histogram of correlation values within and between groups for each reference sequence. The program saves matrices containing raw correlation analysis output in addition to a summary table organized by reference sequence. For phylogenetic comparisons, the program outputs a PHYLIP-formatted file (Fig. 2C) for each reference sequence that summarizes variants within profiles as molecular-morphological characters.

**Figure 2.**
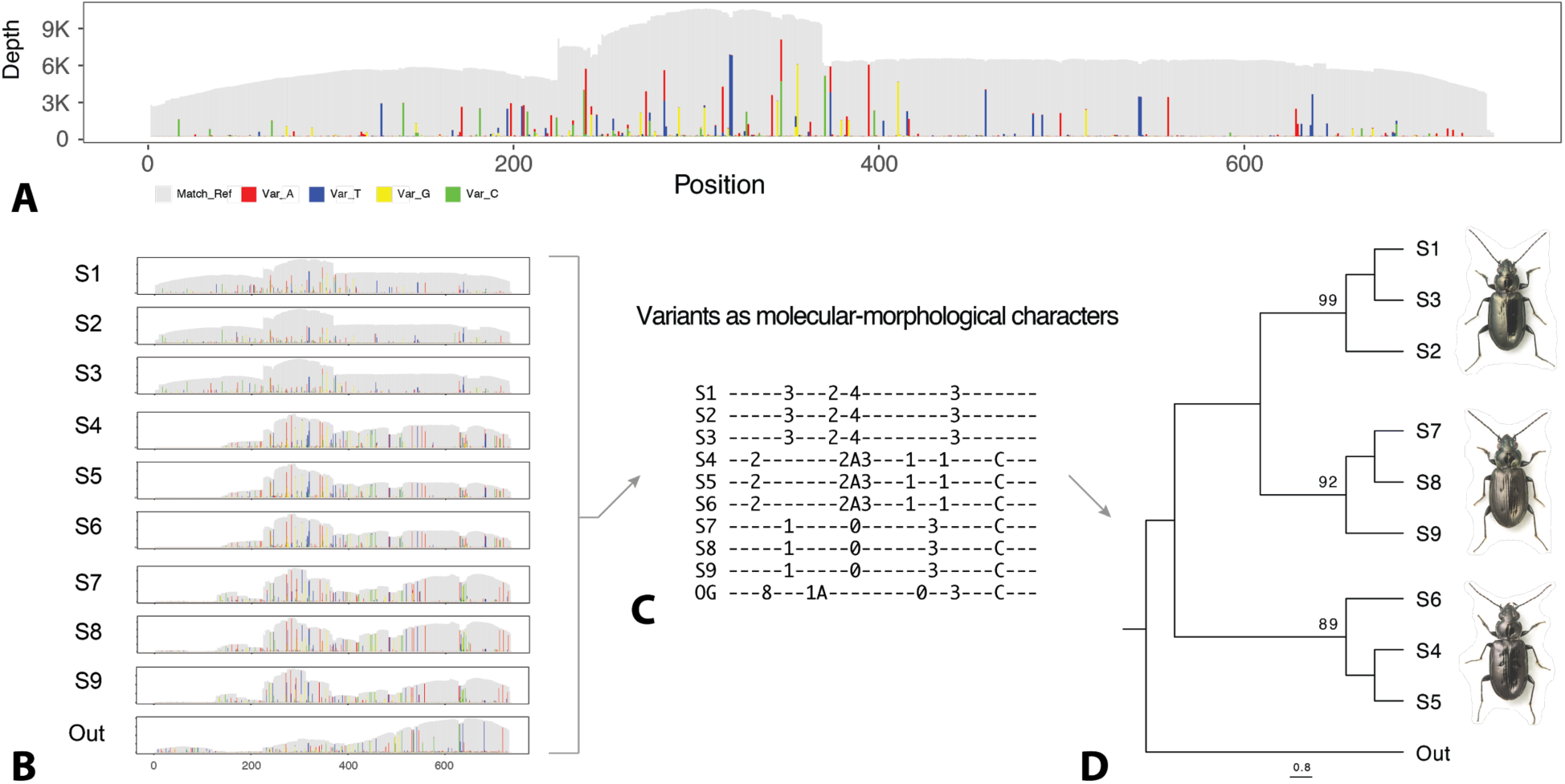
Examples of *RepeatProfiler* variant-enhanced profiles and phylogenetic analysis output. (A) A variant-enhanced repeat profile from the same putative satellite DNA for *B. ampliatum* shown in Fig. 1. (B) Variant-enhanced profiles for the same satDNA in nine individual samples belonging to *B. ampliatum*, *B. breve*, and *B. lividulum*, and a tenth profile for an outgroup *B. aeruginosum* sample. Profiles show species-specific signatures in the pattern of variants relative to the reference sequence. (C) A portion of a PHYLIP file that encodes variant information in profiles as molecular-morphological characters which can be analyzed as using phylogenetic approaches. (D) A tree shows the relationships among samples inferred from variant signals in the profiles in B, produced by analyzing the PHYLIP file produced by the pipeline in IQ-TREE (Nguyen et al., 2015) as morphological data using an MK model.

## Validation results

### Read mapping parameter sensitivity analysis and normalization estimates

The Bowtie2 parameter sensitivity analysis showed only minor differences in the mapping rate of highly conserved reads across Bowtie2 presets ranging from ‘very-fast’ to ‘very-sensitive’, with each run showing ~500X coverage of putatively functional 28S rDNA (Fig. S5). However, parameter settings had a major effect on the mapping rate of divergent 28S-like fragments with the most inclusive setting (*i.e., ‘*-very-sensitive-local’) resulting in a >7-fold coverage increase in mapping of divergent 28S-like reads relative to the least inclusive setting (*i.e., ‘*-very-fast’), and >4-fold increase relative to Bowtie2 default settings (‘-sensitive’) (Fig. S5). Using the ‘-local’ setting as opposed to the Bowtie2 default ‘end-to-end’ setting reduced artefacts of low coverage near the edges of reference sequences (Fig. S6). As a result of these findings, we use ‘-very-sensitive-local’ as *RepeatProfiler’*s default mapping parameters to provide flexibility when mapping reads to non-conspecific reference sequences. In addition, full-length repeats may give rise to various truncated repeats distributed throughout the genome (*i.e.*, small repeats may arise from fragments of larger repeats) as in the example of 28S rDNA in our model species. Thus, permissive default settings allow more layers of repeat evolutionary history to be reflected in profiles. For cases where less permissive mapping is needed, the user can provide their preferred Bowtie2 settings.

Normalization coverage estimates showed a linear relationship with input read number in downsampled data sets well below 1X coverage (Fig. 3A–C). Below ~0.3X coverage, the relationship between reads and average coverage is more variable and leads to a slight over estimation of coverage. This suggests that the normalization feature of the pipeline is robust to low-coverage input reads down to less than 0.5X coverage.

**Figure 3.**
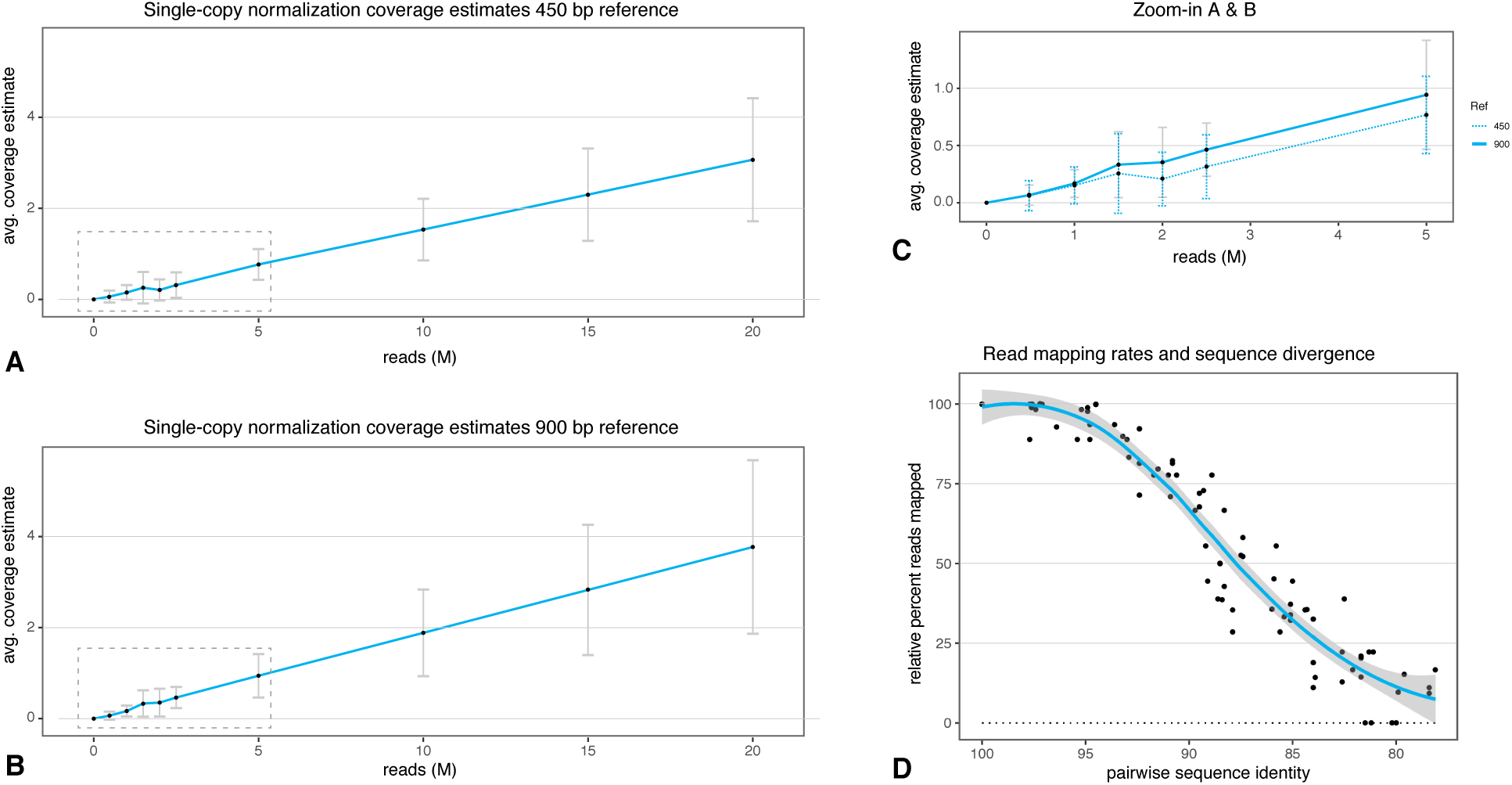
Single-copy normalization estimates and read mapping with sequence divergence. (A) Plot of average coverage estimates obtained by mapping reads to each of ten, 450 bp single-copy reference sequences from *B. lividulum* using the pipeline with datasets ranging from 20 million to 0.5 million reads. The number of input reads shows a linear relationship with avg. coverage estimates down to 0.5 million input reads. The area indicated by the dashed box is shown in ‘C’. (B) Same as ‘A’, but with 900-bp reference sequences. (C) Area shown by dashed boxes in ‘A’ and ‘B’ in greater detail. (D) Results of a simulation study to explore the effect of sequence divergence on read mapping with *RepeatProfiler* default settings. The plot combines data from 80 runs of the pipeline that mapped reads across reference sequences with increasing sequence divergence (0% to 25%) simulated for each of four *Drosophila* species using SLiM 2 (Haller & Messer, 2017).

### Read mapping and reference sequence divergence

Analysis of read mapping trends using simulated data showed no significant difference in mapping rates across the different species and reference sequences analyzed. A plot of the combined data shows that using *RepeatProfiler* default settings, reference sequences with ≤5% sequence divergence show little decrease in mapping rates relative to the unmutated reference sequences (Fig. 3D). Greater than 50% of reads still mapped to references with 13% sequence divergence, and >10% of reads still mapped with 20% divergence (Fig. 3D). These settings allow for considerable divergence between the reference sequence and the sample reads, enabling comparative study across clades spanning shallow to moderate sequence divergence.

### Phylogenetic validation

Phylogenetic analysis of abundant variants in profiles showed that variant signatures can have strong phylogenetic signal over short evolutionary scales. Our in-depth validation using *Drosophila* showed that across 37 TEs from which we generated trees, 28 (75.7%) recovered at least three of the five expected clades with greater than 50% bootstrap support (average bootstrap support = 91.2%) (Figs. S7–8). Of the remaining TEs, five recovered at least two expected clades and four recovered just a single clade. Profiles from 14 TEs recovered all five clades and the correct branching pattern of species. Nine additional TEs produced trees with correct branching patterns, except that one or more replicate samples within species formed a grade rather than clade for that species. We found a single case where a phylogenetic tree had at least moderate support (*i.e.*, greater than 75% bootstrap support) for a relationship in discordance with the species tree (Fig. S7–8). Finding cases where phylogenetic signal strongly contrasts the species tree may be useful for identifying lateral TE transfer among the study taxa.

For 18 of the TEs that underwent phylogenetic analysis, we found data in the literature regarding their recent evolutionary activity (Bergman & Bensasson, 2007). Nine of the 18 TEs are classified as recently active -- these recovered an average of 4.3 of the five expected clades (Table S2). Six of nine recovered all expected clades and the correct branching pattern. The nine inactive TEs recovered an average of 2.4 of the five clades and only one TE recovered all five clades with the correct branching pattern. These findings provide preliminary evidence that, over very short evolutionary time scales, recently active TEs may show stronger phylogenetic signal than TEs whose activity predates speciation events in the study taxa.

## Discussion

*RepeatProfiler* incorporates signals from repetitive sequences into comparative evolutionary studies over short time scales. Unequal crossing over and gene conversion between repeated DNA sequences can cause the concerted evolution of repeats within species and the rapid fixation of differences between species (Coen, Strachan, & Dover, 1982; Gabby Dover, 1994; Gabriel Dover, 1982; Strachan et al., 1982). These rapid evolutionary dynamics can be useful for understanding species boundaries, but repetitive sequences are rarely used in this context. Our approach expands on previous findings (Sproul, Barton, et al., 2020) that repetitive DNA profiles can be stable within, but variable between, closely related species.

The pipeline’s comparative features allow a user to define groups (putative species, populations, male vs female, etc.) *a priori* and test whether signatures in profile shape are correlated with group distinction. In cases where species-specific signatures are not present in profile shape, we show that patterns of sequence variants within profiles may contain such signatures (Figs. S4–S5). Phylogenetic analysis of variant patterns within profiles provides a quantitative approach to group samples based on profile signatures that may also give insights as to phylogenetic relationships of recently diverged taxa (*e.g.*, within species groups) (Figs. 2, 4, S3–4, S6–7), as discussed more below. Combining evidence from many repeats can yield a subset in which recurring patterns become evident. Not all repeats are expected to produce strong signal at the species level, however, the visual enhancements in profiles produced by this pipeline are designed to simplify identification of repeats that show interesting patterns, such as dynamic satellite DNAs and recently active TEs.

**Figure 4.**
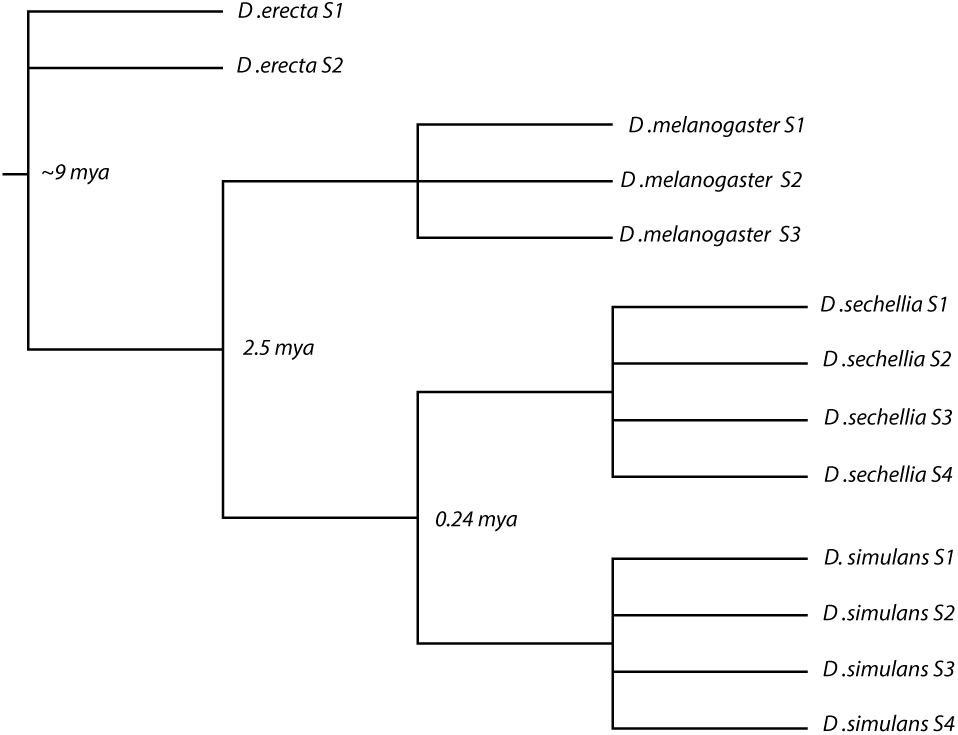
Consensus tree from analysis of profiles for 37 TEs in *Drosophila* species. The consensus tree of all trees generated using PHYLIP files produced by *RepeatProfiler* phylogenetic variant analysis recovered all expected clades with a branching pattern that matches the species tree. The years present of nodes were obtained from (Drosophila 12 Genomes Consortium, 2007; Garrigan et al., 2012).

Previous studies show that patterns of repeat abundance hold phylogenetic signal (Dodsworth et al., 2015); however, repeat abundance can evolve so dynamically (even between sister taxa, let alone across deep phylogenetic splits) that homoplasy can present a major challenge when encoding phylogenetic characters from raw abundance (Martín-Peciña et al., 2019). Rather than use repeat abundance as the character of interest, our approach encodes the signature of abundant variants relative to a common reference sequence. This signature can be highly stable despite fluctuation in absolute copy number that is expected even within species (*e.g*., due to unequal exchange). Our validation in *Drosophila* shows evidence of good phylogenetic signal (Figs. 2, 4, S3–4, S6–7) across divergences as recent as 240 kya (Garrigan et al., 2012), particularly when analyzing recently active TEs. Importantly, we observe this result using reads that have a mixed history of sample preparation and sequencing and a mixture of male and female specimens (Table S1).

In addition to providing high-resolution evidence of species boundaries, repeat profiles can give useful insights into genome architecture and repeat biology. For example, major shifts in repeat abundance across samples can provide evidence of rapid genome evolution through re-patterning of repeat architecture (Sproul, Barton, et al., 2020). Finding uneven coverage of repeats with sharp boundaries can reveal differential amplification of truncated/fragmented copies of that repeat, which suggest the spread of novel satellite DNA sequences (Figs. 1 and S5) or evidence of recent TE activity (Fig. S2). Repeats that show strongly supported phylogenetic relationships that are discordant with species trees can reveal evidence of horizontal transfer. Sex-specific patterns of relative abundance within a species can give insight into dynamics of sex-linked repeats which can be difficult to study with assembly-based approaches, given the highly repetitive nature of sex chromosomes.

When *RepeatProfiler* is used to compare patterns across samples, all sample reads are mapped to a common reference sequence. This approach enables direct comparison of profiles at each position along the reference. Changes observed across samples may be due to both differential abundance of that repeat and/or sequence divergence/indels relative to the reference sequence. The potential limitation of mapping to a common reference sequence is that there is a limit to the extent of divergence which can be included in the same analysis, however, the default parameters of the pipeline enable flexible read mapping across moderate sequence divergence (Fig. 3D).

### *RepeatProfiler* and other programs

Studying repeat dynamics with short-read data has a few distinct advantages, including: (1) the low-cost of data generation; (2) public repositories contain vast amounts of these data from thousands of organisms; and (3) short-read, whole-genome shotgun data approaches can bypass problems caused by gaps and multi-mapping issues in genome assembly-based approaches and allow entire genomic repeat dynamics to be measured (albeit without the context of their position in genomes). designed for studying repeats using short-read data, some of which are outlined in Table 1. We developed *RepeatProfiler* to fill a gap in available tools by allowing efficient study of specific repeats in a quantitative comparative framework.

**Table 1.**
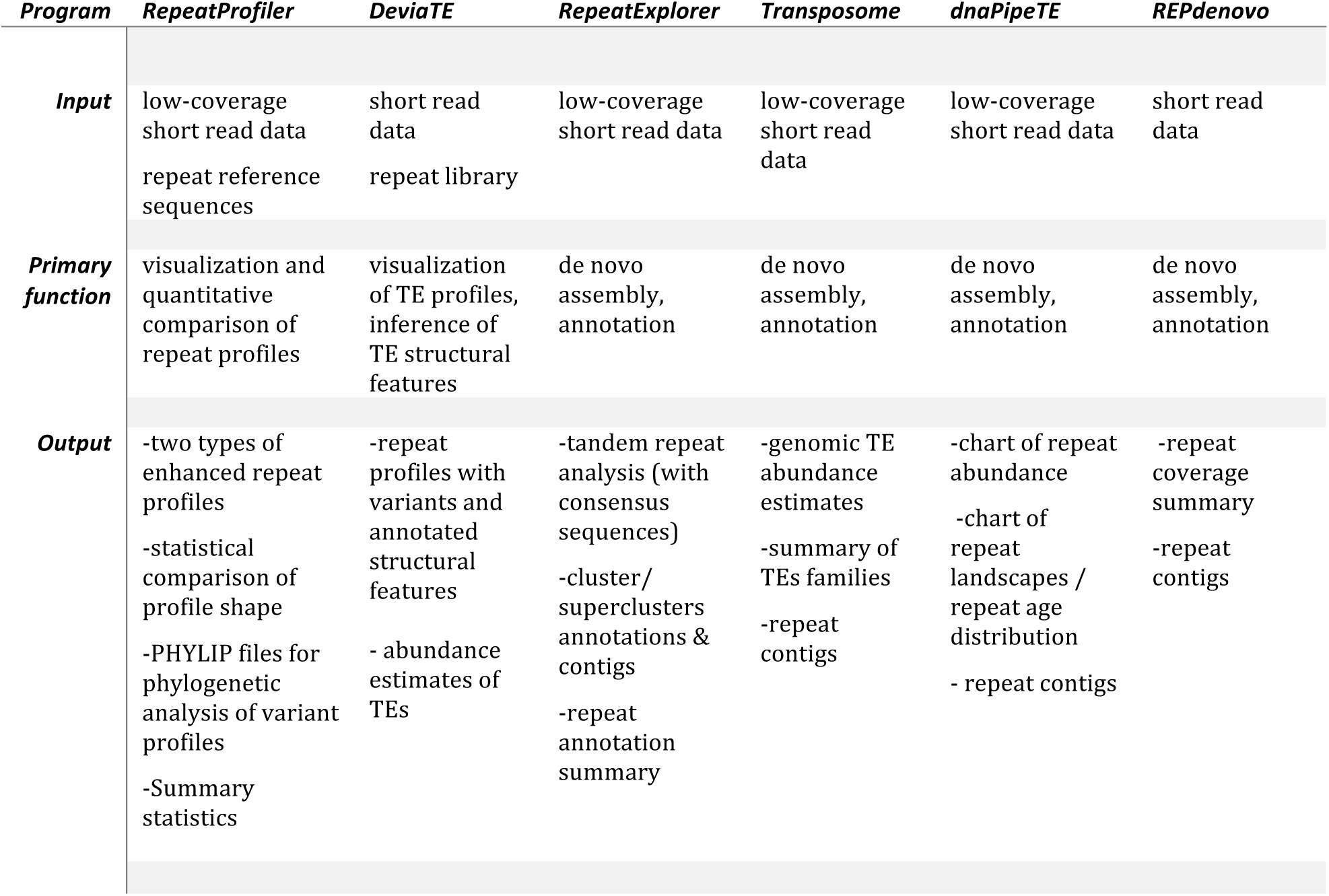
Comparison of RepeatProfiler and other programs for studying repeats with short-read data. This is a list of some existing tools relevant for the present discussion, but is not an exhaustive list of tools nor program functions/output types. Literature cited herein (*e.g*., (Ewing, 2015; Nelson et al., 2017) provides additional summary of available tools.

A growing number of tools are tools exist for *de novo* repeat assembly and initial annotation of repeats from short-read data (*e.g.*, RepeatExplorer (Novák et al., 2013) and dnaPipeTE (Goubert et al., 2015))*. RepeatProfiler* complements these tools, allowing for more in-depth study of specific repeats downstream of those programs. In addition, repeat assembly/annotation programs can be a useful source of reference sequences. For example, RepeatExplorer includes consensus sequences for satellite DNA, LTRs, and rDNA as standard output, which can be directly fed into *RepeatProfiler* as reference sequences.

DeviaTE (Weilguny & Kofler, 2019) is most similar to *RepeatProfiler* in that both tools enable the visualization of read mapping profiles for one or more samples relative to repeat reference sequences. We developed *RepeatProfiler* specifically with the community of researchers studying diverse non-model organisms in mind—its workflow, features, and documentation are designed to support such users in extracting signal from repeats for comparative evolutionary studies in groups with limited resources. Strengths of *RepeatProfiler* include its features that enable quantitative comparison of profile signatures across samples for any repeat type (*i.e.*, satellite DNA, rDNA, TEs, etc.) through correlation analysis of profile shape and through enabling phylogenetic analysis of abundant variants as molecular morphological characters. Although DeviaTE does not enable direct quantitative comparison across profiles, it includes features that offer richer insight into transposable element biology than *RepeatProfiler*. In particular, it can analyze split-reads to detect and annotate specific structural features (*e.g.*, internal deletions, truncations) of transposable elements.

## Acknowledgments

We thank Ching-Ho Chang, Xiaolu Wei, Beatriz Navarro-Domínguez, Lucas Hemmer, Danna Eickbush, Cécile Courret, James Pflug, Antonio Gomez, and Olivia Boyd, and Caitlin Hudecek for helpful feedback on the pipeline and help testing its functionality. This work was supported by National Institutes of Health General Medical Sciences grant R35GM119515, and National Science Foundation grant MCB-1844693 awarded to AML. JSS is supported by an NSF Postdoctoral Research Fellowship in Biology (DBI-1811930).

## Data Accessibility

*RepeatProfiler* is available through GitHub (https://github.com/johnssproul/RepeatProfiler). The data used to demonstrate and validate the tool were downloaded from NCBI’s Sequence Read Archive with accession numbers provided in Table S1.

## Author Contributions

JSS originally conceived the pipeline and its core features. JSS, AML, and SN improved conceptual design and features. SN, JSS, and AG wrote the code. SN, JSS, and AG conducted validation experiments. SN wrote the first draft of the manuscript and all authors contributed to subsequent drafts.

## Notes

https://github.com/johnssproul/RepeatProfiler

### Competing Interest Statement

The authors have declared no competing interest.

