## Supplemental Material for "RepeatProfiler: a pipeline for visualization and comparative analysis of repetitive DNA profiles"

### Supplementary Methods

#### Encoding variant signatures as molecular-morphological characters

For runs that include reads from multiple samples, the pipeline searches for abundant variants relative to the reference sequence and encodes those variants as molecular-morphological characters which can be analyzed with phylogenetic software. Because all samples within a run are mapped to a common set of reference sequences, the relative position of variants can be compared directly across samples for a given reference. For each site along the reference sequence the pipeline records the presence of abundant variants and outputs a PHYLIP-formatted text file with variants encoded as an alpha-numeric character (*e.g.,* an 'a' variant at a given site is encoded '0', 'T' is encoded as '1'). For sites that have multiple abundant variants, that site is encoded with a unique character that represents both variants such that each unique combination of variants is treated as a unique character (*e.g*., a site with 'a' and 't' would be encoded as '5'; 'a' and 'c' is encoded as '6'). The default threshold the pipeline uses to identify 'abundant' variants is 10% of the coverage at a given site. The threshold can be increased or decreased by manually editing a the script that produces the PHYLIP file (see 'percentagecutoff <- 0.1' in scripts/encode_var.R). We routinely analyze the raw output directly in IQ-TREE (Nguyen, Schmidt, von Haeseler, & Minh, 2015) as morphological data with an MK model. In theory the PHYLIP files can be analyzed in any phylogenetic software that can handle morphological data, however some modification of the character coding may be needed to meet formatting requirements of the software. We reiterate here that our approach to analyzing signatures in variant profiles with phylogenetic software does not have a goal of inferring phylogenetic relationships among samples per-se, but rather provides a statistically robust way to extract additional information from repeat profiles to test patterns of sample groupings by evaluating clades within resulting trees.

#### Obtaining repeat reference sequences

Reference sequences may be obtained from existing repeat libraries, online databases, or de novo assembly of repeats from the same low-coverage reads used as input for *RepeatProfiler*. The last source of reference sequences may be the only option for those wanting to explore repeat profiles from low-coverage shotgun reads data in groups with limited genomic resources. This can be done by first characterizing repetitive sequences *de novo* using one of several reference-free assembly/annotation software programs that use short-read data (some are outlined in main text Table 1). To obtain profiles for the putative satellite DNA (satDNA) from main text Fig. 1, we used RepeatExplorer2 (Novák, Neumann, Pech, Steinhaisl, & Macas, 2013) to assemble repeats from low-coverage reads from our beetle data set. We downsampled reads to >1X coverage using seqtk (https://github.com/lh3/seqtk) and uploaded reads to the RepeatProfiler2 using the Galaxy portal (https://repeatexplorer-elixir.cerit-sc.cz) and ran RepeatExplorer2 clustering function. The standard output includes a TAREAN (Tandem Repeat Analyzer in RepeatExplorer) (Novák et al., 2017) analysis, the output for which includes files of consensus sequences of satDNAs, ribosomal DNA, long-terminal repeats, all of which can be downloaded in FASTA format. These can be directly fed into *RepeatProfiler* as repeat reference sequences.

### Supplementary Figures

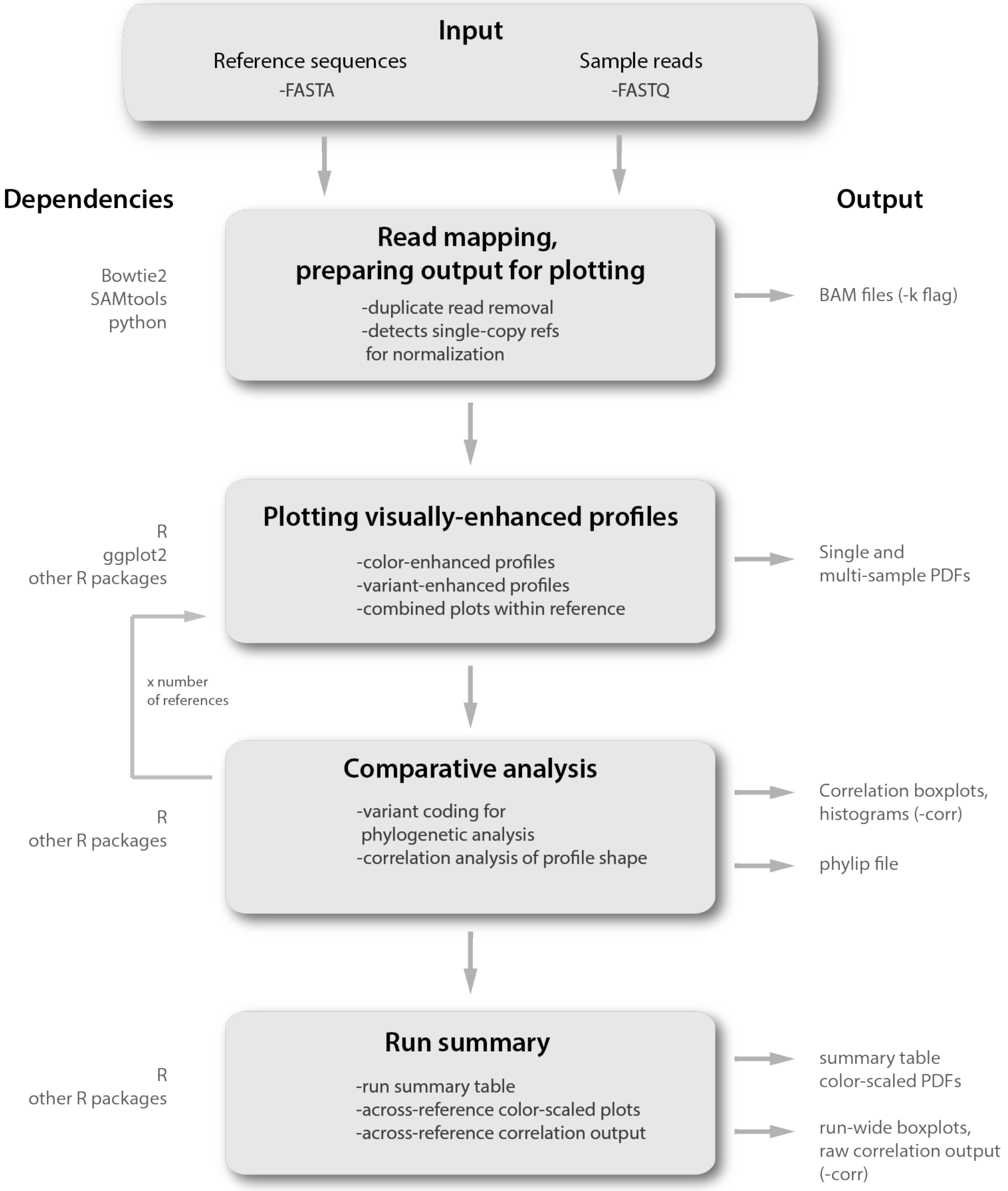


**Figure S1. RepeatProfiler workflow diagram**. Boxes indicate major steps in the pipeline shown in chronological order with dependencies and output products on the left and right, respectively.

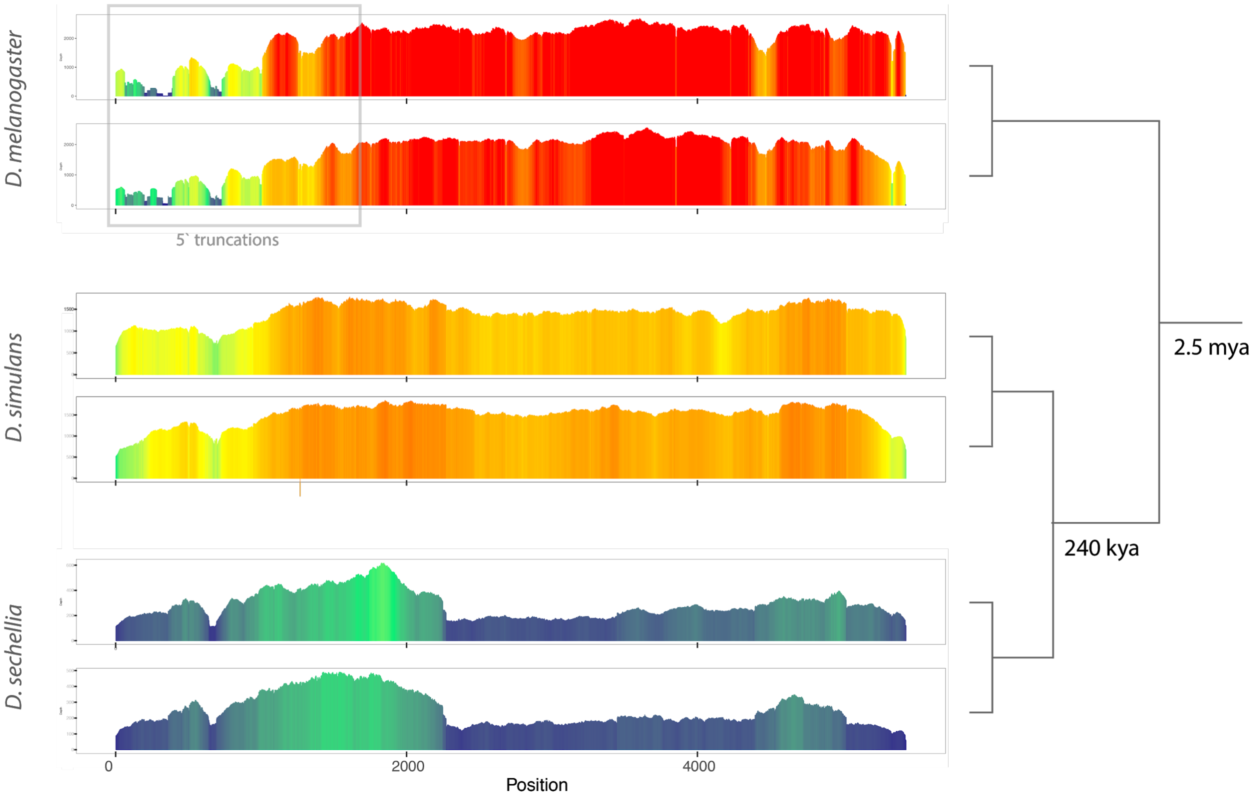


**Figure S2. R1 retrotransposon analysis showing species boundaries and repeat biology.** Profiles of R1 in two individuals for each of three *Drosophila* species show strong species-specific signatures despite their recent evolutionary divergence (*e.g.*, 240 kya -- see tree on right). In addition to providing evidence of species boundaries, repeat profiles can hold information related to repeat biology. Non-LTR elements such as R1 are known to accumulate 5` truncations due to the polymerase falling off during reverse transcription. The pattern of reduced 5` coverage (gray box in *D. melanogaster*) with an otherwise intact element suggests R1 has been recently active in this lineage (consistent with previous (Bergman & Bensasson, 2007).

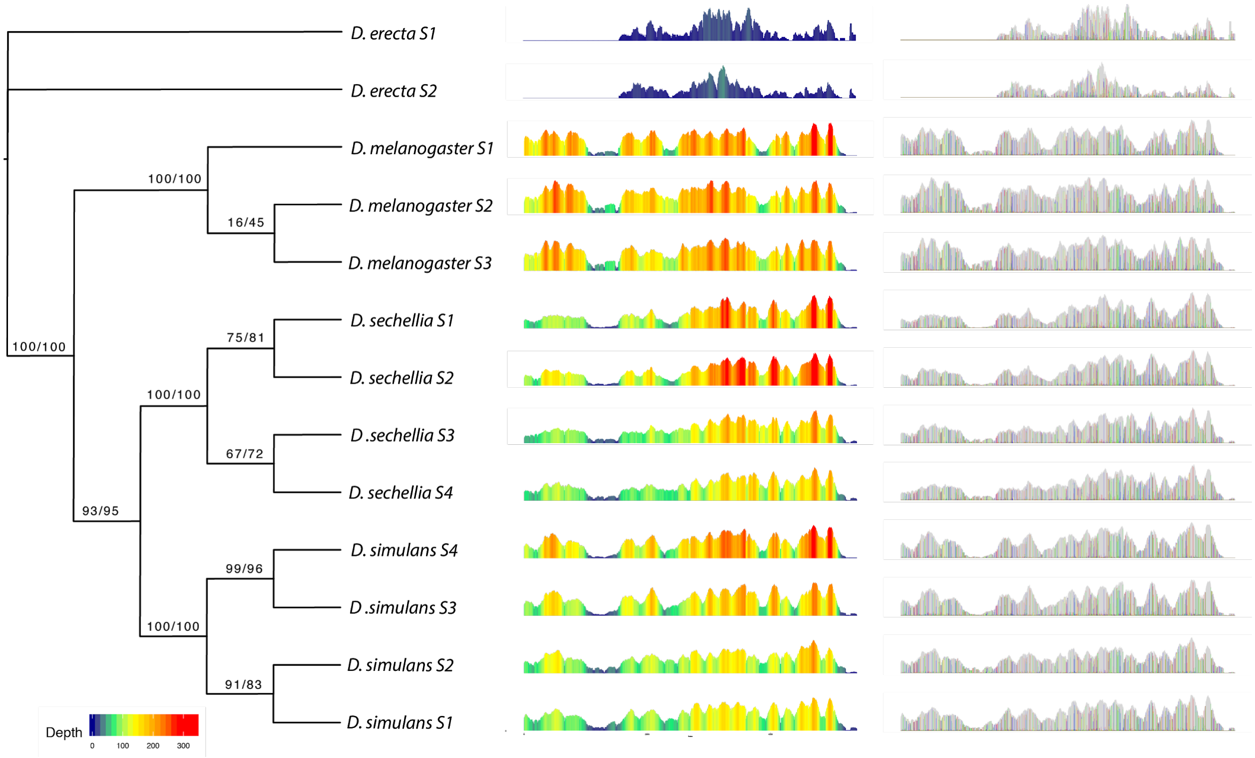


**Figure S3. Phylogenetic analysis of variants can add resolution beyond profile shape**. Color-enhanced profiles for a non-LTR retrotransposon (I-6_DY) in *Drosophila* illustrates an example where profile shape shows limited species-specific signatures between some species (*e.g.,* compare the four *D. sechellia* color profiles to *D. simulans* profiles), however phylogenetic analysis of variant signatures within variant profiles accurately groups conspecific samples with high support. Tree inferred in IQ-TREE with data treated as morphological characters using an MK model.

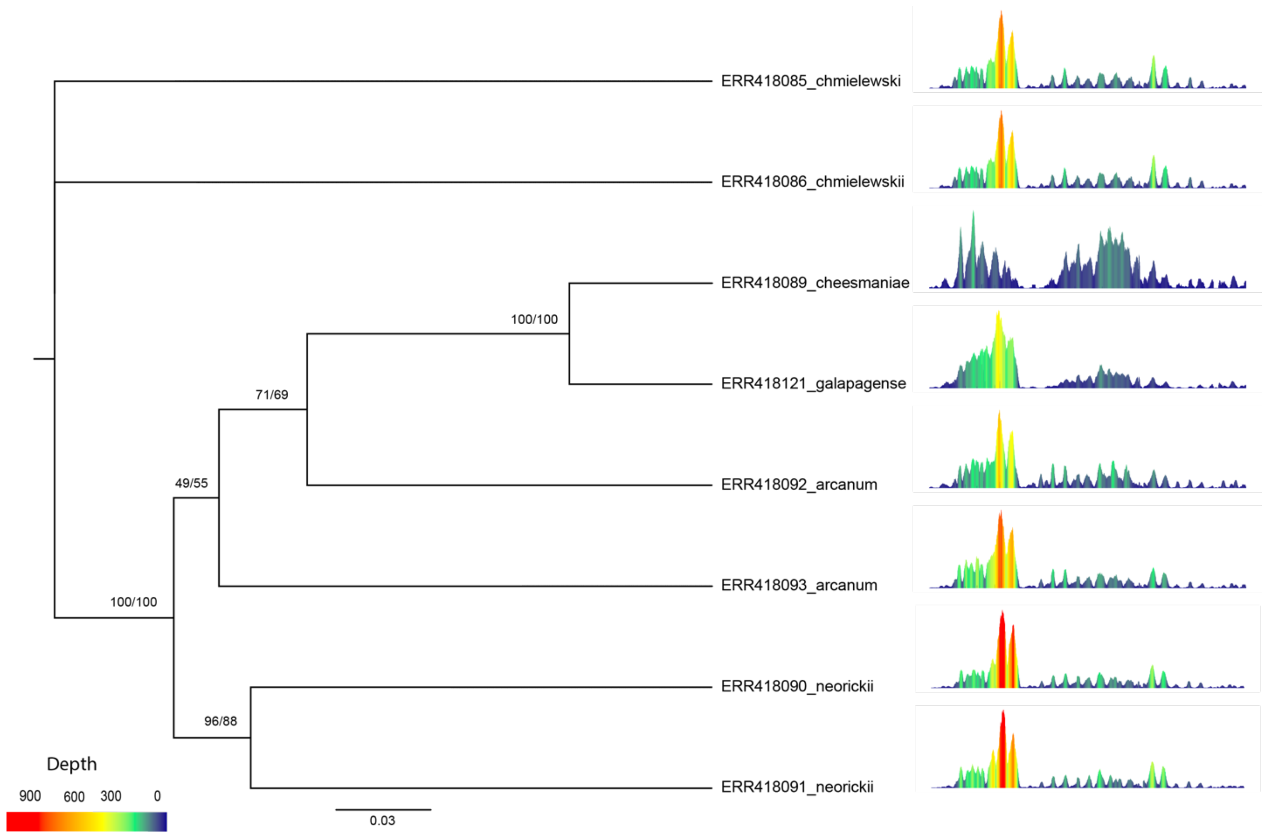


**Figure S4. Phylogenetic analysis of variants in tomato LTRs.** Profiles from an RLX LTR retrotransposons generated from tomato samples representing multiple species shows another example of phylogenetic analysis of variant-enhanced profiles that show signal despite limited signal in profile shape among species**.** Phylogenetic analysis of variant signatures within profiles correctly grouped closely related *Solanum galapagense* and *S. cheesmaniae* as sister (Dodsworth, Chase, Särkinen, Knapp, & Leitch, 2016; Peralta, Spooner, & Knapp, 2008) despite the differences in their profiles. The remaining sample groupings (including the lack of monophyly of the two *S. arcanum* samples) is consistent with relationships inferred from phylogenetic analysis of whole genomes (100 Tomato Genome Sequencing Consortium et al., 2014).

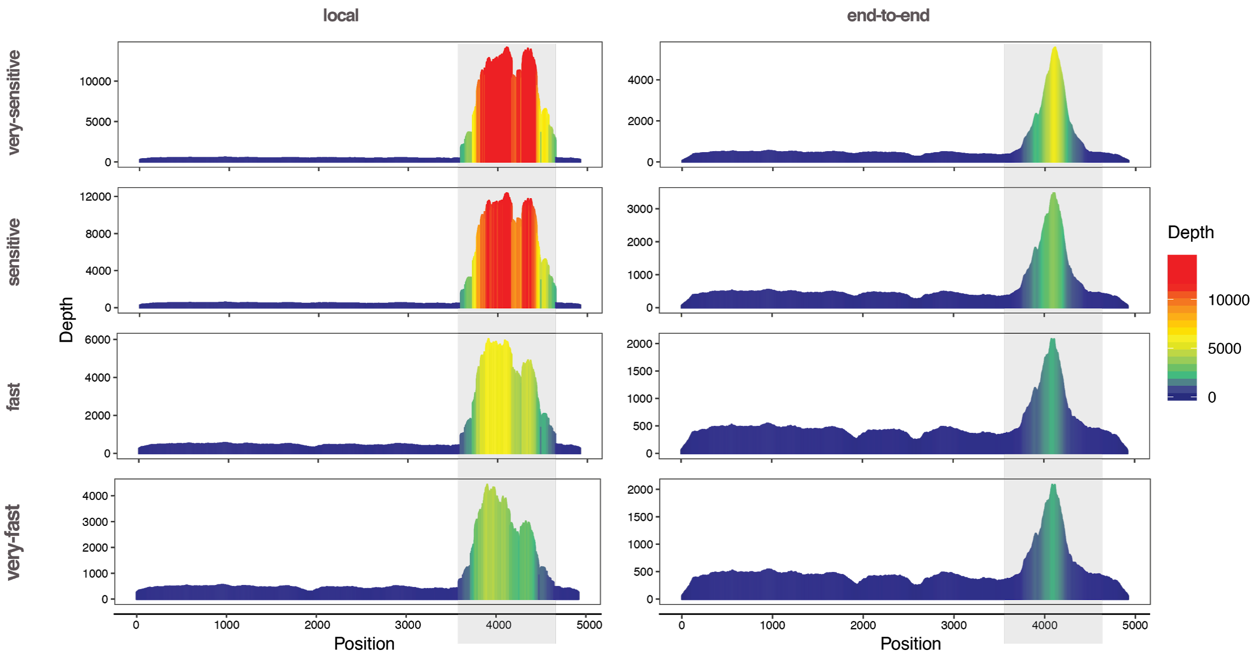


**Figure S5. Bowtie2 parameter sensitivity analysis.** Comparison of profiles generated using Bowtie2's presets: 'very-fast', 'fast', 'sensitive', 'very-sensitive', with 'end-to-end' (Bowtie 2's default) and 'local' mapping strategies. Analysis was conducted with *Bembidion breve* reads, using the 28S ribosomal RNA gene as a reference sequence. This species has a recent history of ribosomal DNA (rDNA) mobilization in which a fragment of 28S rDNA escaped functional rDNA units, proliferated, and spread throughout the genome where it evolves separately (i.e., not in concert with functional rDNA) (Sproul, Barton, & Maddison, 2020). Thus, this species and reference sequence pair is ideal for understanding the effect of changing mapping parameters on both highly conserved (*i.e.,* functional 28S rDNA), and more divergent (abundant rDNA-like fragments) reads relative to the same reference sequence.

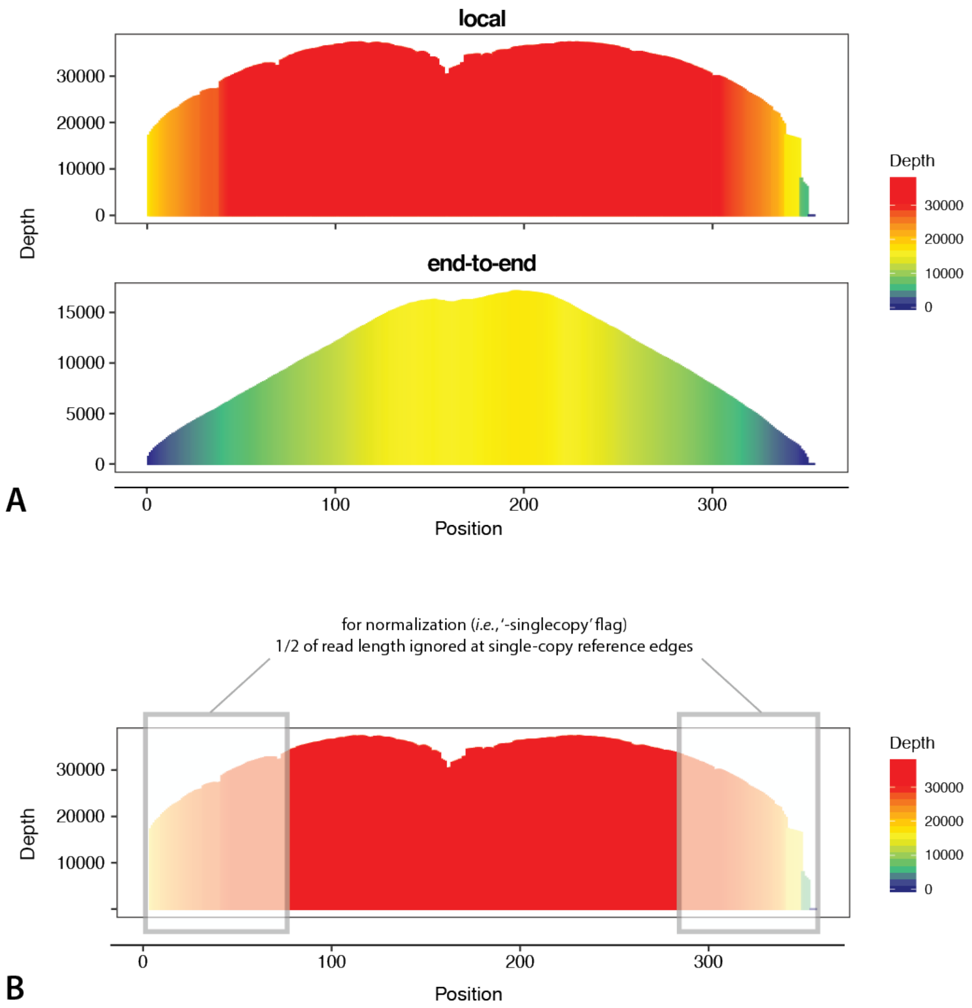


**Figure S6. Correcting edge effects read mapping artefacts.** *RepeatProfiler* takes two approaches to reduce artefacts that are expected near the edges of profiles due to mapping algorithms. (A) The pipeline uses the 'local' mapping setting, as opposed to Bowtie2's default 'end-to-end' setting to reduce artefacts near reference edges that are particularly pronounced on short reference sequences. (B) During normalization to single-copy genes (*i.e.,* the '-singlecopy' flag) the pipeline ignores bases within one half of a read length of the reference sequence edge. For example if reads being mapped have a length of 150 bases then 75 bases will be ignored at each end of the reference in calculating average coverage of single-copy genes (*i.e.,* the gray boxes). This correction reduces the tendency to underestimate average coverage of single-copy genes which would in turn lead to over-estimation of repeat copy number.

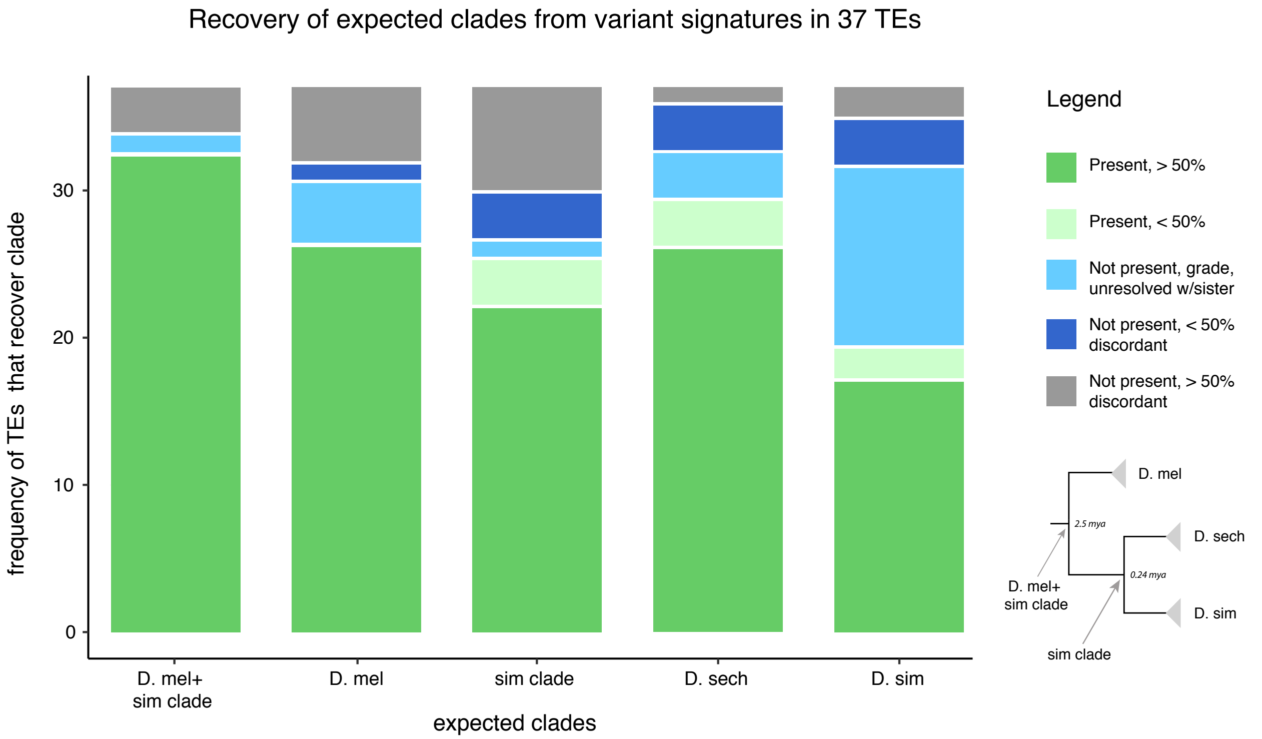


**Figure S7. Recovery of expected clades from variant signatures in 37 TEs.** The x-axis shows each of five clades (see tree schematic in the legend) expected to be present in phylogenetic analysis of variant signatures extracted from profile. Dark green indicates the number of TEs for which the expected clade with greater than 50% bootstrap support in the inferred tree. Light green indicates the number of TEs for which the expected clade was present with less than 50% bootstrap support. Light blue indicates the number of TEs in which the clade was not present due to one or more samples forming a grade (as opposed to a clade), or due to one or more samples sim clade samples showing lack of resolution with its sister taxon. Dark blue indicates the clade was not present with less than 50% support for the discordant (i.e., grouping of non-sister taxa) relationship. Gray indicates the clade was not present with greater than 50% support for a discordant relationship.

| Type | Superfamily | Family | Mel+ simclade | Simclade | Mel | Sech | Sim | Rel | Pseudo |
|---|---|---|---|---|---|---|---|---|---|
| **LTR** | | | | | | | | | |
| | Gypsy | Gypsy_I_LTR_Gypsy | 100 | 100 | 98 | 100 | 93 | X | - |
| | | Gypsy_6B_LTR_Gypsy | 100 | 100 | 99 | 100 | 90 | X | - |
| | | Gypsy-11_DSim-I_LTR_Gypsy | 100 | 100 | 100 | 99 | 89 | X | - |
| | | ZAM_I_LTR_Gypsy | 100 | 95 | 94 | 100 | 99 | X | - |
| | | Gypsy8_I_LTR_Gypsy | 100 | 100 | 78 | 98 | 96 | X | - |
| | | GTWIN_I_LTR_Gypsy | 100 | 95 | 100 | 100 | 76 | X | N |
| | | IDEFIX_I_LTR_Gypsy | 100 | 100 | 100 | 100 | 68 | X | N |
| | | MDG1_I_LTR_Gypsy | 100 | 54 | 98 | 80 | 99 | X | N |
| | | HMSBEAGLE_I_LTR_Gypsy | 100 | 96 | -42 | 100 | 93 | - | N |
| | | Gypsy-8_DSim-I_LTR_Gypsy | 100 | 100 | 94 | 87 | -33 | - | - |
| | | Gypsy2-I_DM_LTR_Gypsy | 100 | 98 | 84 | 91 | -98 | - | - |
| | | STALKER4_I_LTR_Gypsy | 100 | -60 | 60 | 100 | 100 | - | - |
| | | ROVER-I_DM_LTR_Gypsy | 100 | 100 | 89 | 67 | -25 | - | Y |
| | | BURDOCK_I_LTR_Gypsy | 100 | 37 | 67 | 98 | 34 | X | Y |
| | | TIRANT_I_LTR_Gypsy | 62 | 40 | -78 | 94 | 88 | - | Y |
| | | Gypsy5_I_LTR_Gypsy | 100 | -59 | 100 | 100 | -100 | - | - |
| | | DM297_I_LTR_Gypsy | 100 | -50 | 96 | 94 | -50 | - | N |
| | | QUASIMODO2-I_DM_LTR_Gypsy | 100 | -46 | 83 | 43 | -100 | - | N |
| | | Gypsy4_I_LTR_Gypsy | -73 | 83 | 100 | 83 | -83 | - | - |
| | | Invader6_I_LTR_Gypsy | 79 | 79 | -22 | 57 | -30 | - | - |
| | | DM176_I_LTR_Gypsy | 100 | -88 | 85 | -56 | 79 | - | Y |
| | | Gypsy12_I_LTR_Gypsy | -63 | 72 | -63 | -94 | 95 | - | - |
| | | MDG3_I_LTR_Gypsy | -56 | 42 | -56 | 90 | -62 | - | Y |
| | | DM412_LTR_Gypsy | 94 | -61 | -61 | 88 | -61 | - | Y |
| | | NOMAD_I_LTR_Gypsy | 100 | -48 | -48 | -48 | -16 | - | - |
| | | TABOR_I_LTR_Gypsy | 99 | -72 | -72 | -42 | -72 | - | Y |
| | Pao | MAX_I_LTR_Pao | 100 | 100 | -100 | 100 | 78 | - | - |
| | | ROO_I_LTR_Pao | 86 | 80 | 64 | -80 | -59 | - | Y |
| | Copia | Copia1-I_DM_LTR_Copia | 81 | -49 | 83 | -49 | -28 | - | Y |
| **Non-LTR** | | | | | | | | | |
| | R-element | R1_DSe_Non-LTR_retrotransposon_R1 | 100 | 68 | 100 | 86 | 89 | X | N |
| | | R1_DSi_Non-LTR_retrotransposon_R1 | 97 | 81 | 100 | 100 | 55 | X | N |
| | | R2_DM_Non-LTR_retrotransposon_R2 | -11 | -52 | 74 | 42 | -20 | - | - |
| | Jockey | TAHRE_Non-LTR_retrotransposon_Jockey | 74 | 98 | 57 | 100 | 83 | X | - |
| | | HETA_Non-LTR_retrotransposon_Jockey | 95 | 100 | 77 | 100 | -100 | - | - |
| | | DOC_Non-LTR_retrotransposon_Jockey | 81 | -62 | -35 | -62 | -62 | - | - |
| | | FW_DM_Non-LTR_retrotransposon_Jockey | 99 | 59 | 90 | 49 | 29 | X | - |
| | | I-6_DY_Non-LTR_retrotransposon_I | 100 | 98 | 100 | 100 | 97 | X | N |

**Figure S8. Heatmap of phylogenetic recovery of expected clades from profile signatures of 37 TEs.** This figure provides a more detailed summary of Fig. S7 including data from all TEs, while using the same color scheme (see Fig. S7 legend). Numbers in colored cells indicate bootstrap support for the expected clade in green-shaded cells, and support against the expected clade in blue-shaded or gray cells. TEs with an 'X' in the 'Rel' column are cases where all expected clades were present and show a branching pattern that matches the species tree. 'Pseudo' indicates whether TEs were shown to evolve like 'pseudogenes' in (Bergman & Bensasson, 2007). Those not evolving as pseudogenes (*i.e.*, 'N') are expected to be recently active in the history of the group. Those with 'Y' are presumed to be recently inactive.

### Supplementary Tables

**Table STX1. Data used in validation.** This table shows the source of the short-read data we used in our validation experiments in the manuscript.

| Species | Read SRA | Sex | PCR-free | Source |
|---|---|---|---|---|
| *Drosophila* | | | | |
| *D. erecta* S1 | SRR6399451 | Female | Yes | (Wei et al, 2018) |
| *D. erecta* S2 | SRR6399450 | Male | Yes | (Wei et al, 2018) |
| *D. melanogaster* S1 | PRJNA596448 | Female | No | (Sproul et al (2020) |
| *D. melanogaster* S2 | SRR6399449 | Female | Yes | (Wei et al, 2018) |
| *D. melanogaster* S3 | SRR6399448 | Male | Yes | (Wei et al, 2018) |
| *D. melanogaster* S4 | SRR8182349 | NA | No | (Shuhua Fu et al, 2019) |
| *D. sechellia* S1 | PRJNA596448 | Female | No | (Sproul et al (2020) |
| *D. sechellia* S2 | SRR6426002 | Male | No | (Sarmashghi et al, 2019) |
| *D. sechellia* S3 | SRR6399452 | Male | Yes | (Wei et al, 2018) |
| *D. sechellia* S4 | SRR6399453 | Female | Yes | (Wei et al, 2018) |
| *D. simulans* S1 | SRR6399446 | Male | Yes | (Wei et al, 2018) |
| *D. simulans* S2 | SRR6399447 | Female | Yes | (Wei et al, 2018) |
| *D. simulans* S3 | SRR6425999 | Male | No | (Rachtman et al, |
| *D. simulans* S4 | PRJNA596448 | Female | No | (Sproul et al, 2020) |
| *D. mauritiana* S1 | PRJNA596448 | Female | No | (Sproul et al, 2020) |
| *D. mauritiana* S2 | SRR1560267 | Female | No | (Garrigan et al, 2014) |
| *D. mauritiana* S3 | SRR6425993 | Male | No | (Miller et al, 2018) |
| *D. mauritiana* S4 | SRR8834569 | NA | No | (Meany et al, 2019) |
| *Bembidion* | | | | |
| *B. ampliatum* S1 | SRR8530139 | Male | No | (Sproul et al, 2020) |
| *B. ampliatum* S2 | SRR8530136 | Male | No | (Sproul et al, 2020) |
| *B. ampliatum* S3 | SRR8530137 | Male | No | (Sproul et al, 2020) |
| *B. breve* S1 | SRR8530130 | Male | No | (Sproul et al, 2020) |
| *B. breve* S2 | SRR8530064 | Male | No | (Sproul et al, 2020) |
| *B. breve* S3 | SRR8530065 | Male | No | (Sproul et al, 2020) |
| *B. lividulum* S1 | SRR8530146 | Male | No | (Sproul et al, 2020) |
| *B. lividulum* S2 | SRR8530120 | Male | No | (Sproul et al, 2020) |
| *B. lividulum* S3 | SRR8530126 | Male | No | (Sproul et al, 2020) |
| *B. aeruginosum* out | SRR8530105 | Male | No | (Sproul et al, 2020) |
| *Solanum* | | | | |
| *S. chmielewski* S1 | ERR418085 | NA | No | (Schijlen et al.2014) |
| *S. chmielewski* S2 | ERR418086 | NA | No | (Schijlen et al.2014) |
| *S. cheesmaniae* | ERR418089 | NA | No | (Schijlen et al.2014) |
| *S. galapagense* | ERR418121 | NA | No | (Schijlen et al.2014) |
| *S. arcanum* S1 | ERR418092 | NA | No | (Schijlen et al.2014) |
| *S. arcanum* S2 | ERR418093 | NA | No | (Schijlen et al.2014) |
| *S. neorickii* S1 | ERR418090 | NA | No | (Schijlen et al.2014) |
| *S. neorickii* S2 | ERR418091 | NA | No | (Schijlen et al.2014) |
